# Proteomic interrogation of the pathogen-host interface in cholera

**DOI:** 10.1101/2021.01.05.425471

**Authors:** Abdelrahim Zoued, Hailong Zhang, Ting Zhang, Rachel T. Giorgio, Carole J. Kuehl, Bolutife Fakoya, Brandon Sit, Matthew K. Waldor

## Abstract

The microbial cell surface is a critical site of microbe-host interactions that often control infection outcomes. Here, using the infant rabbit model of cholera, which provides an abundant source of *in vivo Vibrio cholerae* cells and diarrheal fluid, we investigated the proteomic composition of this interface. Bulk diarrheal fluid proteomes revealed that cholera toxin accounts for the vast majority of the host proteins present during infection. We developed a surface biotinylation protocol to purify and quantify both bacterial and host proteins present on the surface of diarrheal fluid-derived *V. cholerae*. We found that SP-D, a toxin-dependent host protein that directly binds the *V. cholerae* surface, is a novel intestinal defense factor. Other *V. cholerae*-bound host proteins also bound distinct taxa of the murine intestinal microbiota. Proteomic investigation of the microbial surface-host interface should be a valuable tool for probing microbe-host interactions and their influence on homeostasis and infection.

## Introduction

The principal interaction site between microbes and their hosts is found at the microbial cell surface. Microbial surface proteins, such as pili, adhesins, and receptors often mediate direct interactions with host cells as well as the host milieu, facilitating microbial growth (1–3). Conversely, host-derived proteins such as antimicrobial peptides, antibodies, and complement bind to the bacterial surface and often restrict microbe growth (1). To date, comprehensive characterization of the bacterial cell surface proteome or the host proteins that bind the pathogen during infection has been challenging, due in part to the technical constraint of obtaining sufficient quantities of *in vivo* pathogen cells for analysis. However, recent advances in tandem mass tag (TMT) mass spectrometry (TMT-MS), and the availability of cell-impermeable protein labeling reagents (4–9), suggest that the development of new approaches to monitor the proteome of the microbial surface *in vivo*, including at the pathogen-host interface during infection, should be feasible.

The rod-shaped, Gram-negative bacterium *Vibrio cholerae* causes the severe and potentially lethal diarrheal disease cholera, which remains a significant threat to global public health. Cholera has afflicted humans for centuries, remains endemic in over 50 countries (10) and has caused major recent outbreaks, e.g. in Haiti and Yemen (11,12). Cholera is caused by consumption of food or water containing *V. cholerae*. The hallmark symptom of cholera is large quantities (up to 20L/day in severe cases) of watery diarrhea, which can contain up to 10^9^ cfu/ml of *V. cholerae* (13). The production of choleric diarrhea is thought to promote the pathogen’s dissemination in the environment and subsequent transmission to naïve hosts.

Studies in human volunteers have established that choleric diarrhea is caused by the activity of cholera toxin (CT), an AB_5_-type toxin that is secreted by *V. cholerae* in the small intestine (10,14–16). The enzymatic A subunit of CT catalyzes the ADP-ribosylation and constitutive activation of G_s_ alpha subunits within small intestinal epithelial cells, increasing the activity of adenylate cyclase and leading to elevated intracellular cAMP concentrations, which in turn stimulate active efflux of sodium (Na^+^), chloride (Cl^−^), potassium (K^+^), bicarbonate (HCO ^−^), and water out of the cells (10,17). Orogastric inoculation of purified CT is sufficient to trigger cholera-like diarrhea in humans (18). Human studies have also revealed the importance of the toxin co-regulated pilus (TCP), a *V. cholerae* cell surface appendage whose expression *in vivo* is activated by the same virulence regulatory network as CT, and is essential for colonization of the small intestine (19). Additional surface-associated factors, e.g. heme transport proteins and outer membrane proteins (OMPs) also facilitate *V. cholerae* survival and growth in the small intestine (20–22).

Cholera is restricted to humans, but several animal models have been developed to study *V. cholerae* intestinal colonization and diarrheal disease (23). Much has been learned regarding *V. cholerae in vivo* biology and pathogenicity from suckling (~3-day-old) rabbits, where orogastric inoculation with *V. cholerae* leads to robust intestinal colonization and a disease that closely mimics severe human cholera (24). As in humans (19), *V. cholerae* intestinal colonization is TCP-dependent in this model. Moreover, infant rabbits develop large volumes of CT-dependent watery diarrheal fluid, and CT is also sufficient to induce diarrhea in these animals (24). In infant rabbits, diarrheal fluid accumulates to high levels in the cecum (approximately 0.5-1mL/animal) prior to excretion. The fluid contains a high density (10^9^-10^10^ cfu) of *V. cholerae,* and thus provides a relatively pure source of *in vivo* organisms that has been leveraged for various high-throughput investigations, including RNA-seq analyses of the pathogen’s *in vivo* transcriptome (25) and Tn-Seq analyses of its genetic requirements for *in vivo* growth (26–29).

Limited analyses of the *V. cholerae* proteome *in vivo* (30,31) have been reported. Previous efforts using activity-based protein profiling (ABBP) defined the active serine hydrolases in diarrheal fluid of infant rabbits (32). This study suggested that secreted *V. cholera* proteases, including IvaP, decreased the activity of host proteases in diarrheal fluid (32,33). This work also suggested that the amount of intelectin, a host intestinal lectin bound to *V. cholerae in vivo* was reduced by secreted *V. cholerae* proteases. This observation raised the intriguing possibility that the pathogen is bound and/or targeted by a previously undefined set of host proteins as it transits through the gastrointestinal tract. However, studies to define the diarrheal fluid proteome and the role of *V. cholerae* factors in triggering the release of host proteins have not been reported.

Here, we used the infant rabbit model of cholera and TMT-MS to define the bulk proteome of choleric diarrhea. Additional TMT-MS analyses of surface labelled diarrheal fluid-derived *V. cholerae* cells (in an approach termed Surface Protein LAbelingS Host/Microbe, SPLASH/M) enabled identification of both pathogen and host proteins present at this interface. Unexpectedly, we discovered that CT accounts for nearly the complete set of >1000 proteins identified in diarrheal fluid. We found that one of the most abundant CT-dependent proteins, surfactant protein D (SP-D), directly binds *V. cholerae* and functions as a region-specific intestinal defense factor. SPLASH/M identified the suite of *in vivo V. cholerae* cell surface proteins and also revealed a number of host-derived bacterial-binding proteins (HBBP), that were not previously known to interact with bacteria. In addition to SP-D and Intelectin, these HBBPs include Lactoperoxidase, Annexin A1 and Zinc-Alpha-Glycoprotein. Notably, we found that these proteins could not only associate with *V. cholerae,* but with the surfaces of a subsets of murine gut symbiotic bacteria, suggesting that HBBPs may facilitate intestinal bacterial homeostasis. The SPLASH/M approach provides a new lens to reveal the pathogen-host interface and should be applicable to define the microbe-host proteomic interface in a wide variety of settings.

## Results

### Cholera toxin drives the host proteomic response to *Vibrio cholerae*

To investigate how CT impacts the host proteomic response to *V. cholerae*, we used the infant rabbit model of cholera. The chemistry of the diarrheal fluid that accumulates in the cecum in this model resembles that of choleric fluid (24), but its proteomic composition has been less characterized. In particular, while CT is known to induce secretion of Cl^−^ and water into the intestinal lumen, the pathogen factors that lead to the accumulation of proteins in choleric fluid are unknown. Infant rabbits were oro-gastrically inoculated with wild-type (WT) *V. cholerae* (an isolate from the Haiti 2010 outbreak (34)), a derivative of the WT strain containing a deletion of *ctxAB* (V*. cholerae* Δ*ctx*), or purified CT (50 μg), to assess the contribution of CT in stimulating the proteomic response to *V. cholerae* intestinal colonization (Fig. 1A). Mock infected rabbits that were inoculated with buffer only served as a negative control in these experiments. Consistent with previous reports (35,36), the burden of WT and V*. cholerae* Δ*ctx* in the diarrheal fluid were similar, but there was much greater abundance of diarrheal fluid in the animals that were inoculated with WT versus Δ*ctx V. cholerae* (Fig. S1B). There was at least as much cecal fluid recovered from animals inoculated with CT alone as animals inoculated with WT *V. cholerae*, supporting the idea that CT is the major determinant of fluid accumulation in this model. There was a small amount of diarrheal fluid obtained from mock infected animals that was sufficient for proteomic analysis.

**Figure 1:**
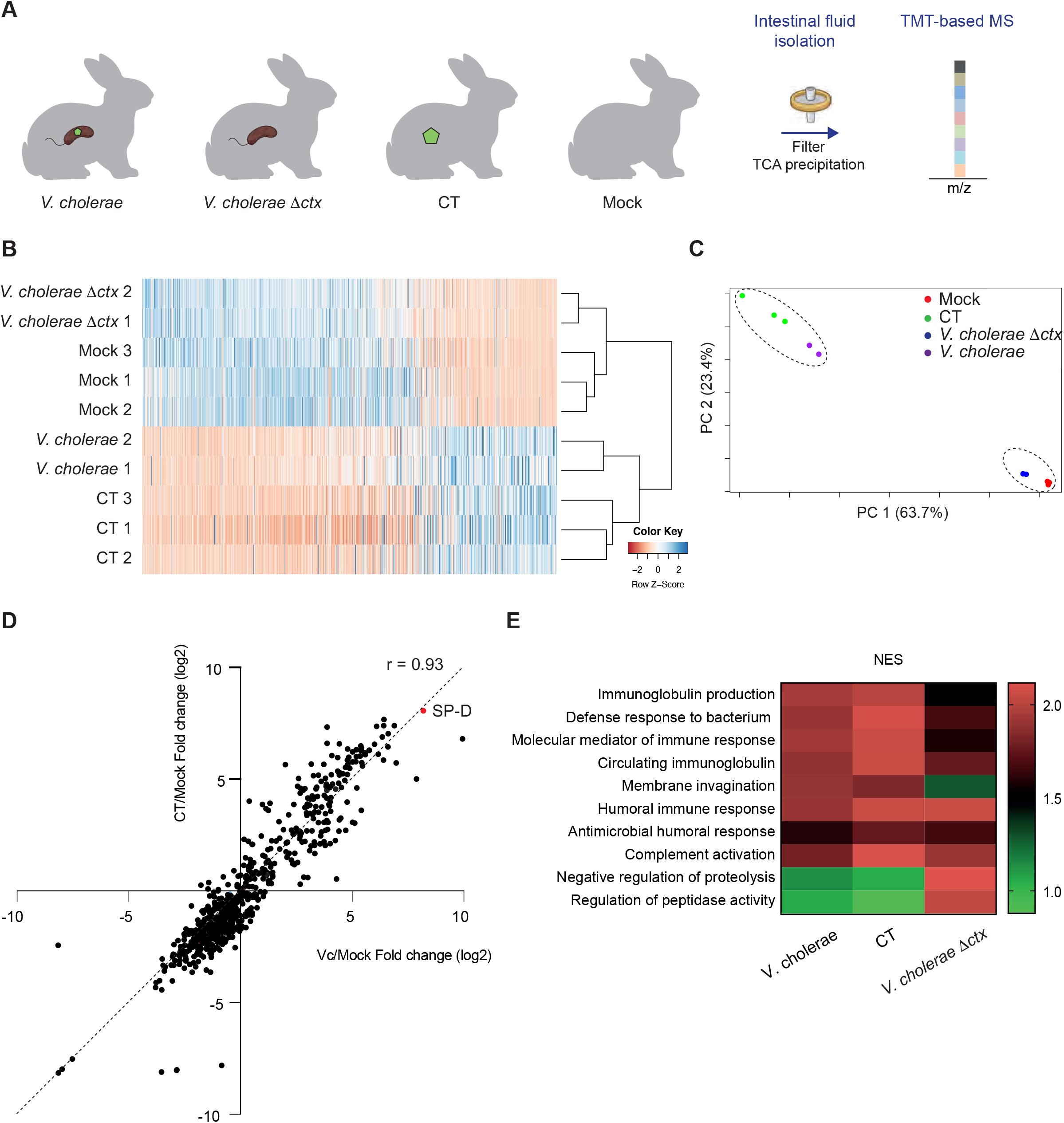
The diarrheal fluid proteome in infant rabbits is largely stimulated by CT. (A) Schematic of the experimental protocol for identification of the proteome in diarrheal fluid isolated from rabbits inoculated with *V. cholerae*, *V. cholerae* Δ*ctx*, purified cholera toxin (CT) or buffer alone (Mock). (B-C) Hierarchical clustering and principal component analysis (PCA) of proteomes identified by TMT-based mass spectrometry. (B) Heatmap is sorted by the log2 fold change between WT *V. cholerae* infected and mock. (D) Scatterplot showing relative fold changes in the abundance of proteins isolated from rabbits inoculated with either wild-type *V. cholerae* (Vc) or purified cholera toxin (CT), each relative to the proteomes of control animals. The red dot indicates SP-D. (E) Comparison of the gene sets enrichment from the GO molecular function pathways for the proteomes of rabbits infected with *V. cholerae*, *V. cholerae* Δ*ctx*, or CT infected rabbit versus control animals. NES: normalized enrichment score. Pathways were considered to be significantly enriched if the adjusted p-value was less than 0.25.

High-resolution tandem mass tag (TMT) mass spectrometry (4) was used to quantitatively analyze the protein composition of the diarrheal fluid isolated from the four groups of rabbits. The 5968 peptides identified in this analysis were mapped to the rabbit proteome (37) and corresponded to 1014 different proteins including 664 identified with more than 1 peptide (Fig. 1B, Table S1). Most of the proteins were predicted to be extracellular (Fig. S1C), consistent with the idea that *V. cholerae* intestinal colonization does not disrupt the integrity of the intestinal epithelial barrier (10,15) and lead to the release of cytoplasmic proteins into the intestinal lumen. Unexpectedly, both unsupervised hierarchical clustering and principal component analysis revealed that the protein composition of fluid from animals infected with *V. cholerae* Δ*ctx* was very similar to that in control animals, suggesting that in the absence of CT, *V. cholerae* intestinal colonization does little to alter the secretion/release of host proteins into the intestinal lumen (Fig. 1B-C). These analyses also revealed that the protein composition of fluid from animals infected with WT *V. cholerae* or treated with CT alone were very similar (Fig. 1B-C-D). Relative fold changes in individual protein abundance in samples from animals given CT only or infected with WT *V. cholerae* were strongly correlated (r = 0.93), further underscoring the similarity of the proteomic signatures of these fluids (Fig. 1D, S1D). Together, these observations strongly suggest that the activity of CT, in addition to triggering the secretory response of ion and water flow into the intestinal lumen, also drives the secretion/release of hundreds of proteins that are found in the cholera-like diarrheal fluid of infant rabbits.

Pathway enrichment analysis revealed several pathways specifically associated with WT *V. cholerae* infection and CT treatment (Fig 1E, Table S2). These included several GO Biological Process terms linked to immune responses, including “immunoglobulin production” and “defense response to bacterium”, suggesting that CT plays a role in modulation of the immune response to *V. cholerae*. WT *V. cholerae* infection and CT treatment also led to similar reductions in relative abundances of proteins classified as regulators of proteolysis, raising the possibility that CT modifies host protease activity.

### SP-D directly binds *V. cholerae* and is an intestinal mucosal defense factor

One of the most abundant proteins in diarrheal fluid samples from animals infected with WT *V. cholerae* or given CT was surfactant protein D (SP-D). SP-D is a C-type lectin that mediates pulmonary innate immune defense and has been recently reported to function in intestinal homeostasis by impacting the composition of the gut microbiota (38–41). The proteomic data suggested that SP-D was ~60-fold enriched in the CT-only and WT *V. cholerae* infections compared to the mock infected controls (Fig. 1D, red dot). Western blotting of filtered diarrheal fluid from uninfected and WT-infected rabbits with a polyclonal anti-SP-D antibody confirmed that rabbit SP-D was highly enriched in diarrheal fluid from infected rabbits (Fig. 2A)

**Figure 2:**
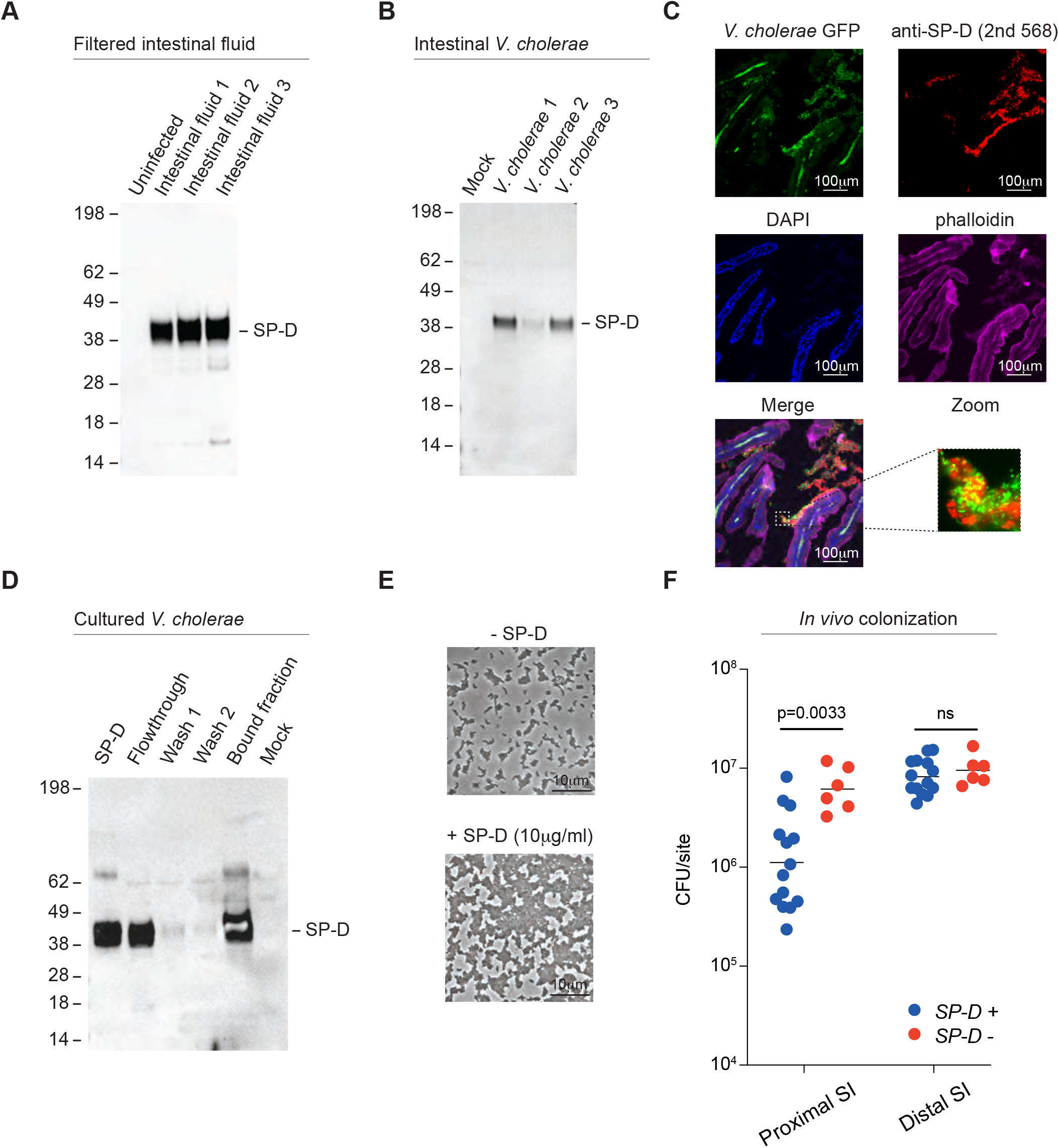
SP-D is an intestinal mucosal defense factor. (A) Detection of SP-D in filtered diarrheal fluid by immunoblotting. Diarrheal fluids were collected, filtered and TCA precipitated before western blotting using anti-SP-D antibody. (B) Detection of SP-D associated with *V. cholerae* cells isolated from diarrheal fluid. (C) Immunofluorescence micrographs of rabbit small intestines inoculated with *V. cholerae*-GFP. Bacterial cells were detected by GFP fluorescence, SP-D was detected with a goat anti-SP-D antibody followed by anti-goat antibody coupled to Alexa fluor 468. Phalloidin (for actin labeling) is stained with an antibody coupled to Alexa fluor 647 and DAPI (for DNA labeling) is shown is blue. Scale bar is 100 μm. (D) Immunoblot detection of recombinant SP-D incubated with *V. cholerae* cells grown in LB. From left to right: purified SP-D protein, flowthrough (unbound protein), washes and bound fraction were analyzed alongside with bacterial cells treated with buffer only (mock). (E) SP-D aggregates *V. cholerae* cells. Bacterial cells were incubated in PBS containing 5mM CaCl_2_ for 1 hour in the presence (lower panel) or absence (upper panel) of SP-D (10 μg/ml) and analyzed by light microscopy. Scale bar is 10 μm. Results from an experiment representative of three independent experiments are shown. (F) *V. cholerae* small intestinal colonization in littermate *sftpd^−^/^−^* and *sftpd^+^/^+^* mice. Bacterial burdens recovered from proximal and distal small intestine 18 hrs after *V. cholerae* inoculation. Note S-PD+ include both heterozygotes (*sftpd^+^/^−^*) and homozygous (*sftpd^+^/^+^)* animals.

We wondered whether this CT-induced lectin binds to *V. cholerae* during infection. To test this possibility, *V. cholerae* cells were isolated from the diarrheal fluid of infected infant rabbits, filtered, washed, lysed, and immunoblotted for SP-D. We readily detected SP-D in the *V. cholerae* collected from the diarrheal fluid, suggesting that SP-D associates with *V. cholerae* cells *in vivo* (Fig. 2B). Immunofluorescence microscopy was used to further investigate the association of SP-D and *V. cholerae in situ* during infection. In these experiments, infant rabbits were inoculated with a fluorescently-tagged wild-type *V. cholerae* strain (*V. cholerae* GFP) and sections from the small intestines of infected animals were stained with an antibody to SP-D (Fig. 2C, S2A). Both *V. cholerae* and SP-D co-localized to the region immediately above the epithelium (Fig. 2C), whereas staining with the secondary antibody used to detect the antibody to SP-D did not bind to rabbit tissue on its own (Fig. S2A). At higher magnification (Fig. 2C, see zoom), SP-D and *V. cholerae* GFP localize within the same matrix. A similar staining pattern was reported for WGA-positive mucin aggregates in *V. cholerae*-infected rabbits (24), suggesting that SP-D might co-localize with mucin and *V. cholerae* during infection.

SP-D harbors a C-terminal carbohydrate recognition domain that mediates its interactions with microorganisms (38,42,43). We adapted a previously described bacterial whole-cell ‘pull-down’-like assay (32), using purified SP-D protein, to test whether this protein could directly bind *V. cholerae*. In these experiments, *V. cholerae* cells grown *in vitro* were incubated with human SP-D (0.25μg) and then the flow through (unbound), washes, and bound fraction (lysed cells) were analyzed for the presence of SP-D using immunoblots. A band corresponding to SP-D, as detected in the positive control (SP-D lane), was observed in the flow through and bound fractions, whereas almost no SP-D was detected in the two wash fractions. These observations demonstrate that SP-D can directly interact with *V. cholerae* cells in the absence of an intermediary host factor (Fig. 2D).

Some host lectins can promote the agglutination of target cells (42,44,45). For example, SP-D leads to agglutination of the fungal pathogen *Pneumocystis carinii* (*jiroveci*) (38) and the bacterial pathogen *Streptococcus pneumoniae* (42). Incubation of *V. cholerae* cells with human SP-D *in vitro* also led to their agglutination, indicating that SP-D binding can alter *V. cholerae* physiology (Fig. 2E, wide field in Fig. S2B). We next used the well-established suckling mouse model of cholera to investigate if SP-D impacts *V. cholerae* intestinal colonization. For these experiments, heterozygous (*sftpd*^+/−^) breeders were used to generate litters that contained both *sftpd*^+/+^/*sftpd*^−/+^ (SP-D^+^) and *sftpd*^−/−^ (SP-D^−^) offspring. Littermates of suckling mice were inoculated with WT *V. cholerae* and bacterial burdens in the proximal and distal small intestine were enumerated 18 hrs after inoculation (Fig. 2F). There were significantly higher *V. cholerae* burdens in proximal small intestinal samples from SP-D^−^ vs SP-D^+^ mice, suggesting that SP-D contributes to intestinal defense. In contrast, there was no difference in the number of *V. cholerae* recovered from the distal small intestines of SP-D^−^ and SP-D^+^ mice. Thus, SP-D’s protective function appears to be limited to the proximal small intestine.

### Identification of *V. cholerae* and host proteins on the pathogen cell surface during infection

The observation that a host protein SP-D, is bound to the *V. cholerae* cell surface during infection led us to hypothesize that additional host proteins are also present at this pathogen-host interface. To identify these factors, we adapted an unbiased approach that has been used to define how viral or parasitic infection leads to changes in the landscape of the surface proteome of eukaryotic cells (5,6,46–48). To identify both bacterial-and host-derived proteins present at the surface of *V. cholerae* cells collected from infected animals, we developed Surface Protein LAbelingS Host/Microbe (SPLASH/M). In this approach, total surface proteins associated with *V. cholerae* cells isolated from the diarrheal fluid of infected rabbits were first labeled with the cell-impermeable primary amine biotinylation reagent Sulfo-NHS-SS-Biotin (Fig. 3A). After labeling, bacterial outer membrane fractions were purified and subjected to affinity purification to isolate biotinylated proteins. Then, TMT-based mass spectrometry was used to identify and quantify the labelled proteins. The biotinylated fraction contained the known outer membrane protein OmpU, but not the cytoplasmic RNA polymerase subunit RpoB, confirming that the bacterial lysis protocol did not lead to cytoplasmic contamination of the biotinylated fractions (Fig. S3A-C). Critically, host proteins bound to the *V. cholerae* cell surface as well as bacterial surface proteins were labeled using this protocol, and detected peptides were mapped to both the *V. cholerae* and the rabbit genomes.

**Figure 3:**
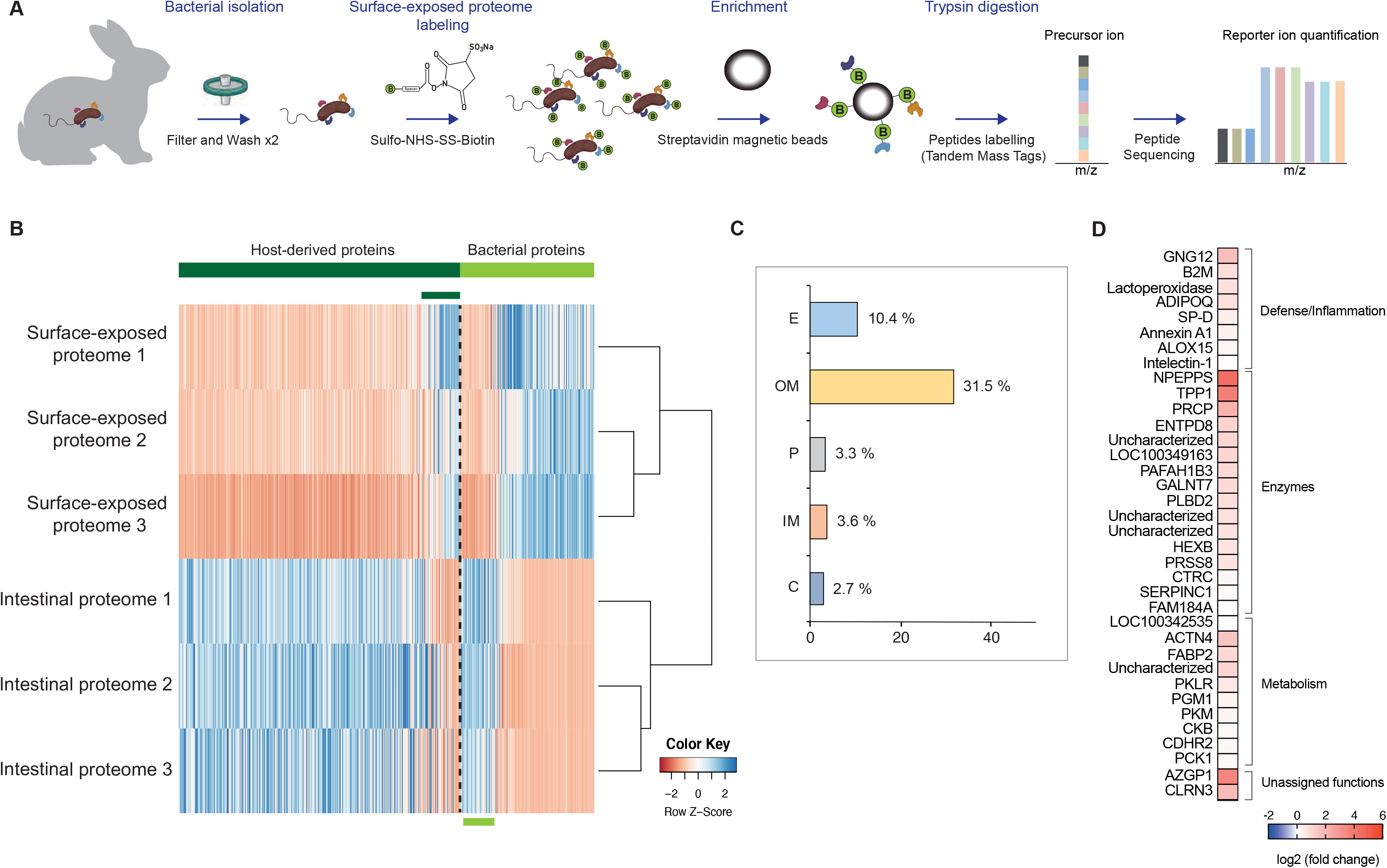
Identification of surface-exposed *V. cholerae* proteins and *V. cholerae*-bound host-derived proteins. (A) Schematic of the SPLASH/M protocol. (B) Hierarchical clustering of the surface proteins identified by SLASH/M and the proteins identified in diarrheal fluid in three animals. Host-derived and bacterial proteins were sorted separately by log2 fold change prior to clustering. (C) Proportion of the proteins identified by SPLASH/M relative to total number of ORFs encoded in the *V. cholerae* genome for each predicted localization (extracellular (E), outer membrane (OM), periplasmic (P), inner membrane (IM) or cytoplasmic (C)). (D) Heat map showing the ratio of abundance of host-binding bacterial proteins (HBBPs) identified with SPLASH/M versus their abundance in the diarrheal fluid of the corresponding animal.

To contextualize the host proteins found in the biotinylated fractions, we also performed TMT-MS analysis on cell-free fractions of the diarrheal fluid, from which bacteria, particulate matter, and host cells had been removed by filtration. After removing proteins for which only 1 peptide was found, we identified 564 total proteins, including 382 rabbit and 182 bacterial proteins, across all conditions (Table S3). Hierarchical clustering revealed that the proteins identified in the three surface labeled samples and the three diarrheal fluid samples clustered together and exhibited distinct proteomic profiles (Fig. 3B). As expected, the abundance of most rabbit proteins was greater in the fluid samples, except for a small subset of ~35 proteins that were more abundant in the surface labeled samples (Fig. 3B). Similarly, a subset of ~50 *V. cholerae* proteins was more abundant in the fluid samples (Fig. 3B). Notably, these proteins included CT, Xds (VC2621), a secreted nuclease involved in mediating escape from neutrophil extracellular traps and in degradation of extracellular DNA (49) and PrtV, a metalloprotease implicated in *V. cholerae* virulence (50). Several *V. cholerae* cell surface-associated virulence factors known to be up-regulated *in vivo* (25), such as TCP components, HutA, an outer membrane heme receptor (51), and the accessory colonization factor AcfA, required for efficient intestinal colonization (52) and present in the outer membrane vesicles (OMVs) *V. cholerae* produces *in vivo* (31) were included in the surface labelled samples, providing further biological validation to the dataset. Bioinformatic predictions of protein subcellular localization revealed that 38% of the surface-labeled *V. cholerae* proteins were outer membrane proteins (Fig. 3C), constituting ~32% of the total predicted *V. cholerae* outer membrane proteome. In contrast, only 2.7% of the total predicted *V. cholerae* cytosolic proteome were labeled, reinforcing the idea that there was minimal cytoplasmic contamination in the labeled samples. CT was one of the surface-labeled proteins, suggesting that a fraction of CT remains associated with the cell surface prior to its secretion. Notably, 46% of the proteins previously identified in *V. cholerae* OMVs released during infection (31) were identified in the surface-labeled proteome (Fig. S3C), consistent with the idea that OMVs contain a subset of surface-associated proteins.

The most intriguing set of proteins identified with SPLASH/M were the 36 rabbit proteins that were enriched on the *V. cholerae* surface compared to diarrheal fluid (Fig. 3D). One of these HBBP was Intelectin, a lectin previously found to be associated with *V. cholerae* during infection (32) and another was SP-D, which was shown to bind *V. cholerae* above (Fig. 2). Some of the HBBPs (8/36), like SP-D, are proteins with known or predicted roles in host defense and inflammation; however, most of these proteins are thought to function in pathways that are not directly related to host defense and are not known to associate with bacterial cells.

### LPO, Annexin A1, and ZAG directly bind *V. cholerae*

Three of the identified HBBPs, Lactoperoxidase (LPO), Annexin A1 (AnxA1), and Zinc-alpha-2-glycoprotein (ZAG, or AZGP1) were chosen for further study. Although each of these proteins has been reported to be in the extracellular space, none have been shown to bind bacteria. LPO and AnxA1 have been implicated in innate defense (53–55). LPO generates the antimicrobial hypothiocyanite in presence of H_2_O_2_, and is expressed in secretions including milk and saliva and on mucosal surfaces including the intestinal epithelium (55,56). Annexin A1 is generally thought of as a host cell surface death marker (57), and *V. cholerae* proteases have been found to modulate its abundance in the intestines of infected rabbits (32). ZAG is a soluble protein present in serum and other body fluids that has been associated with diverse non-immune functions (58,59). Notably, however, ZAG exhibits a major histocompatibility complex (MHC) like-structure and exhibits structural similarities to beta-2-microglobulin (B2M) (60,61).

To corroborate the proteomic data, we first probed *V. cholerae* samples isolated directly from the diarrheal fluid of infected rabbits for the presence of LPO, AnxA1 and ZAG by Western blotting. Bands corresponding to each protein were detected on the bacterial cells collected from infected animals, suggesting that LPO, AnxA1 and ZAG associate with *V. cholerae* cells during infection (Fig. 4A-C).

**Figure 4:**
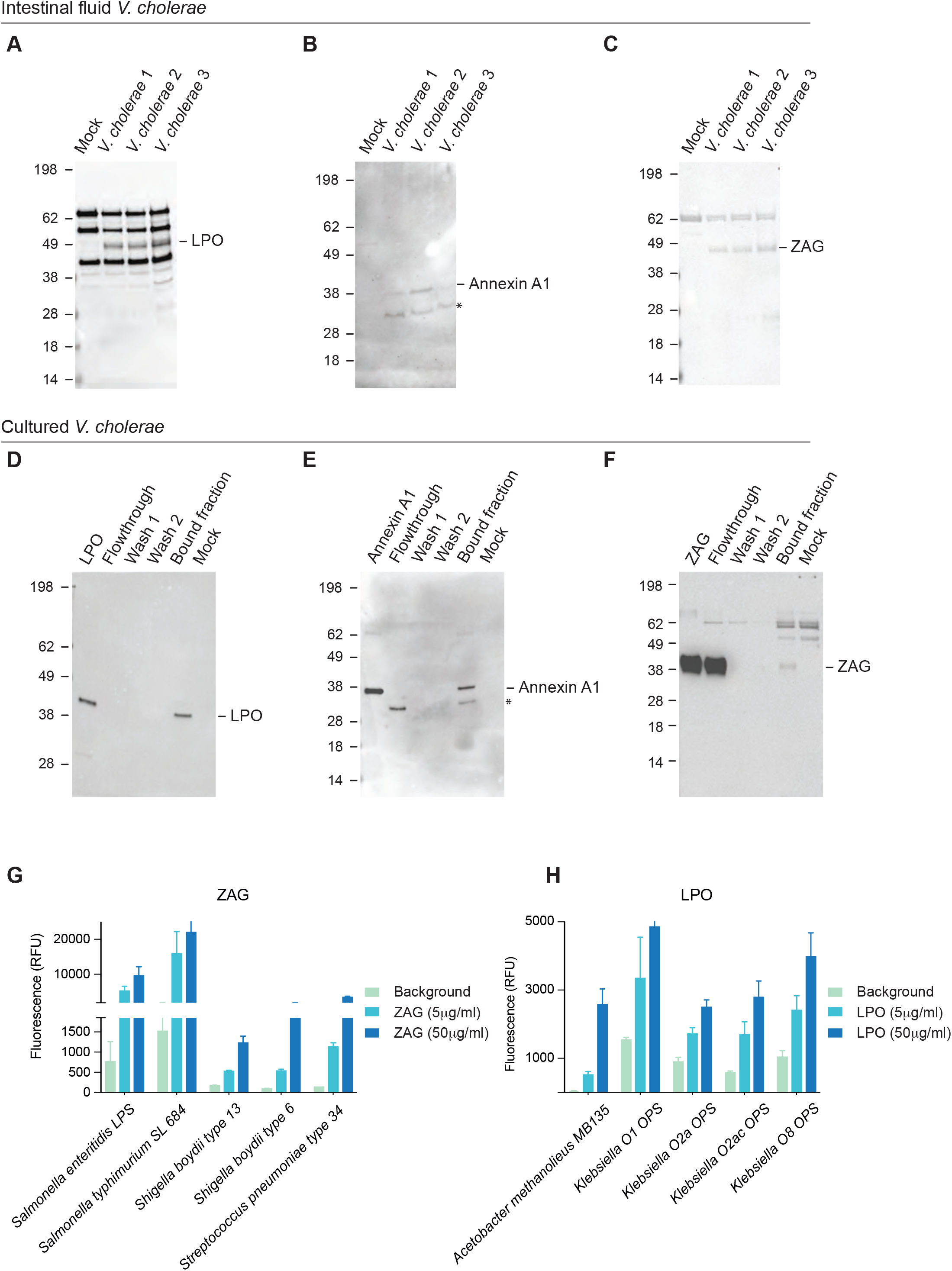
LPO, AnxA1 and ZAG interact with *V. cholerae*. (A-C) Detection of LPO, AnxA1 and ZAG associated with *V. cholerae* cells in the intestine*. V. cholerae* cells collected from diarrheal fluid of infected rabbits were washed twice and lysed. Proteins were separated by 10% acrylamide SDS-PAGE and immunoblots for LPO (A), AnxA1 (B) and ZAG (C) were performed. (D-F) Detection of LPO, AnxA1 and ZAG associated with *V. cholerae* cells grown in the laboratory. Immunoblot detection of recombinant LPO (D), AnxA1 (E) and ZAG (F) incubated with *V. cholerae* cells cultured in LB. From left to right: purified proteins, flowthrough (unbound protein), washes and bound fraction were analyzed alongside with bacterial cells treated with buffer only (mock). (G-H) ZAG and LPO glycan binding assessed by glycan microarrays. Binding of recombinant human ZAG (5 μg/ml and 50 μg/ml) (G) and LPO (5 μg/ml and 50 μg/ml) (H) to microbial glycan arrays. Data are shown as mean ± s.d. (n = 4 technical replicates). (Glycan array data organized by genus are in Supplemental Fig. S4 and the full dataset in Supplemental Table S4)

Next, we tested if these proteins directly interact with *V. cholerae* grown in the laboratory using the binding assay described above (Fig. 4D-F). For all three proteins, a band corresponding to the molecular weight of the respective purified protein was detected in the elution fraction, though the amount of ZAG bound was not as great as the other two proteins. Little or no protein was observed in the wash fractions for any of these proteins, suggesting that each protein can interact with *V. cholerae* in the absence of additional host factors (Fig. 4D-F). Apparent proteolysis of Annexin A1 was detected in the *in vivo* as well as *in vitro* assays consistent with the previous report that *V. cholerae* proteases can cleave Annexin A1 (Fig. 4B, Fig. 4E, see star Ref).

We reasoned that the capacity of LPO, AnxA1 and ZAG to bind microbes was not likely to be restricted to *V. cholerae* and hypothesized that these HBBPs may bind to conserved microbial cell surface structures such as glycans or phospholipids. To test whether these three proteins bind to microbial glycans, we used the glycan microarrays developed by the Consortium for Functional Glycomics (CFG; http://www.functionalglycomics.org/). These microarrays contain more than 300 highly purified and characterized bacterial polysaccharides isolated from a broad range of diverse microbes (62), but do not include *V. cholerae* polysaccharides. In these experiments, two doses (5 and 50μg/ml) of each HBBP were put on the arrays and after incubation, were washed, and binding was detected with a fluorescent secondary antibody; the signal measured with the secondary antibody alone was used to set background levels of detection (Fig. 4GH, and S4 A-B). Both ZAG and LPO exhibited dose-dependent binding signals to different polysaccharides, whereas AnxA1 did not (Fig. 4GH, and S4 A-B). ZAG bound to *Salmonella* and *Shigella boydii* LPS and the capsular polysaccharide from *S. pneumoniae* 34 (Fig. 4G, top 5 hits). LPO bound to polysaccharides from different microbes, including *A. methanolieus*, and *Klebsiella* (Fig. 4H, top 5 hits). Together, these data suggest that although ZAG and LPO are not considered lectins, they can bind to structurally diverse microbial glycans.

### HBBPs interact with gut commensal bacteria

Given the binding of LPO and ZAG to bacterial glycans, we hypothesized that these HBBPs, as well as AnxA1, may also bind to symbiotic organisms within the gut microbiota. Consistent with this idea, a previous study found that SP-D bound to ~2% of fecal bacteria (39). We developed a modified ‘IgA-Seq’-like method (63) to isolate and identify symbiotic microbes that are bound by LPO, AnxA1 or ZAG in the intestine (Fig. 5A). Microorganisms were isolated from the feces of specific-pathogen-free (SPF) mice and labeled with the DNA-specific dye SybrGreen, to facilitate differentiation of living microbes from food debris. HBBP-coated microorganisms were detected by flow cytometry, using biotinylated anti-HBBP antibodies and Cy7-conjugated streptavidin (Fig. 5A). While almost no bacteria were labeled by the Cy7-strepavidin secondary reagent, antibodies to LPO, AnxA1 or ZAG, were found to bind to 1-10% of the fecal microbiota in SPF mice. LPO coated a somewhat higher fraction of microbes (6.5% ± 3) than Annexin A1 (3% ± 2), and ZAG (2% ± 3; Fig. 5A-B). Thus, these 3 HBBP, like IgA and SP-D, interact with gut symbionts.

**Figure 5:**
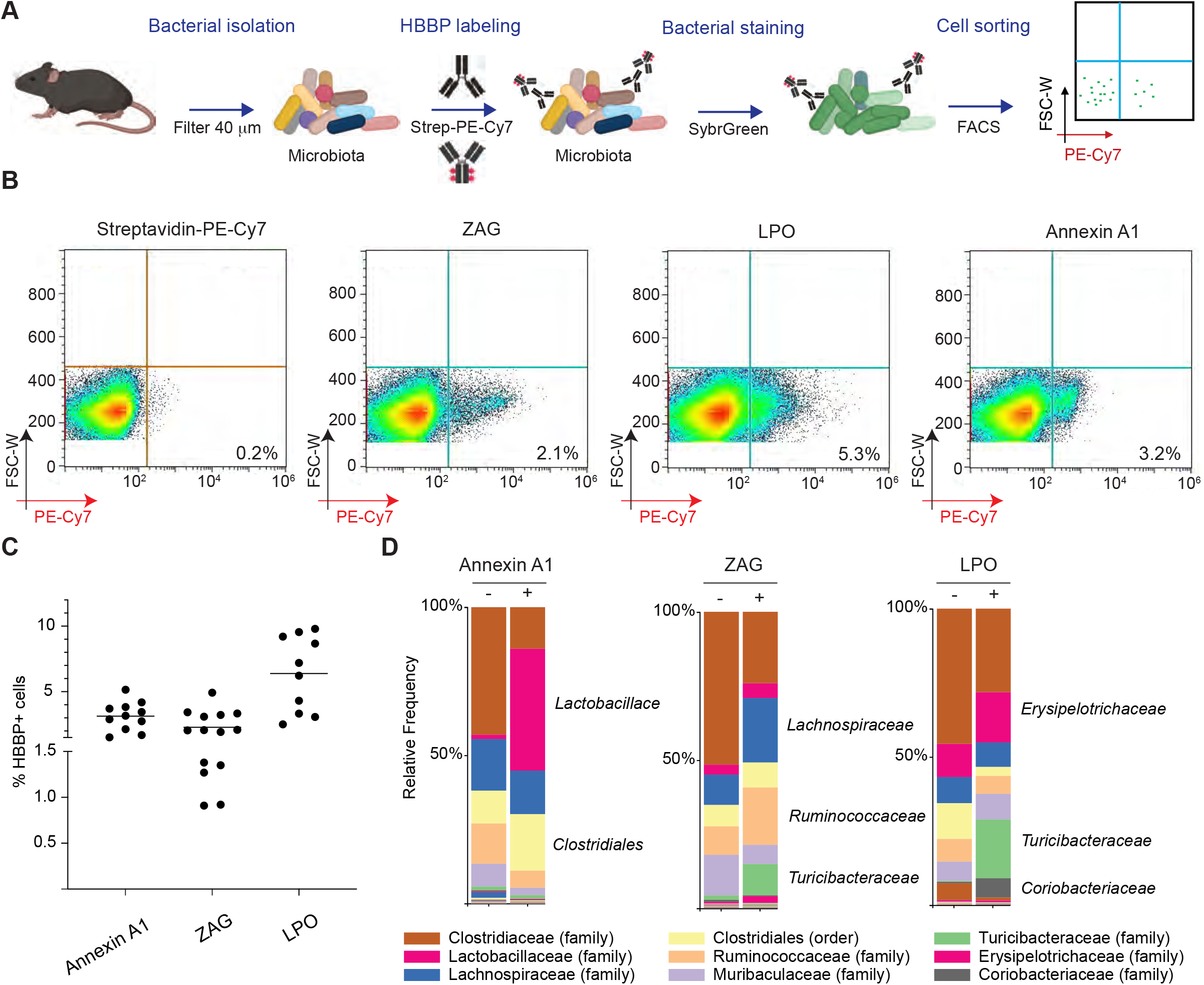
HBBPs interact with gut commensal bacteria. (A) Schematic of the workflow for detection of HBBP bound to fecal microbiota. (B) Flow cytometry of microbiota stained with Streptavidin-PE-Cy7 only, or antibodies to ZAG, LPO or AnxA1. (C) Quantification of flow cytometry data from (B). (D) Relative abundance of order or family-specific taxonomic units (OTUs) after 16s rRNA sequencing of sorted cells from (A). The bound (positive) and unbound (negative) fraction of the microbiota is shown. Each bar represents the average from four individual mice.

To compare the taxonomy of the HBBP-coated vs -uncoated microbial species, we used FACS to sort the HBBP-bound (and hence fluorescently tagged) and unbound bacterial fractions in each sample, and then carried out 16S rRNA sequencing analysis to classify the populations (Fig. 5C). PCA showed that the positive (HBBP-coated) and negative (unbound) populations for each HBBP were distinct (Fig. S5A). The uncoated bacteria from all three analyses generally clustered together, whereas the LPO-, AnxA1-, and ZAG-coated bacteria formed distinct clusters, suggesting that these HBBPs bind to distinct microbial species. Accordingly, operational taxonomic unit (OTU) distributions between coated and uncoated populations differed; the coated bacteria were enriched for different OTUs for all three of these proteins (Fig. 5D). In particular, 1) LPO-coated bacteria were enriched in *Lactobacillaceae*, *Turicibacteraceae* and *Coriobacteriaceae*; 2) ZAG-coated bacteria were enriched in *Lachnospiraceae*, *Ruminococcaceae* and *Turicibacteraceae*; and 3) Annexin A1-coated bacteria were highly enriched in *Lactobacillaceae*. These data suggest that these 3 proteins interact with taxonomically distinct microbes.

## Discussion

Despite more than a century of research on cholera pathogenesis (64) there is limited knowledge of the protein content of choleric diarrheal fluid, and of the proteomic landscape of the *V. cholerae* cell surface-host interface during infection. Using the infant rabbit model of cholera, we unexpectedly discovered that CT, *V. cholerae*’s signature virulence factor, is almost solely responsible for the pathogen’s impact on the host’s secretion or release of proteins during infection. SP-D, one of the most abundant proteins in diarrheal fluid, was found to be associated with the *V*. *cholerae* surface during infection and to impede *V. cholerae* colonization in the proximal small intestine. We developed SPLASH/M, to chart the protein landscape of both bacterial and additional host proteins present at the pathogen surface during infection. 36 host-derived bacterial binding proteins (HBBP) were identified with this approach. Additional studies of three of these proteins, LPO, AnxA1, and ZAG, corroborated their capacity to bind *V. cholerae* and demonstrated that LPO and ZAG can bind to bacterial glycans. Moreover, these proteins were found bound to distinct symbiotic bacteria in the gut microbiota, suggesting that HBBPs may modulate the composition and function of host-associated microbial communities. These observations suggest that approaches to define the proteins present at the microbial surface-host interface, such as SPLASH/M, will provide a valuable new tool for understanding microbe-host interactions.

Cholera toxin’s primary role in *V. cholerae* pathogenesis is generally thought to be in facilitating pathogen dissemination and transmission by dramatically increasing the volume of diarrhea in infected individuals. In experimental animals, CT only has marginal impacts on *V. cholerae* intestinal colonization burden per se (35,65), but since there are massive numbers of the pathogen shed in diarrheal stool, the toxin has a major net impact on pathogen replication and dissemination. Furthermore, recent studies suggest that CT impacts the nutrient composition of the intestinal milieu in infected animals, potentially supporting optimal *in vivo V. cholerae* replication/ colonization. Besides stimulating secretion of Cl^−^ and water into the intestinal lumen, CT is also known to exert additional effects on intestinal epithelial cells, including goblet cell degranulation, leading to mucus secretion (24,66,67). It is possible that at a subset of the ~1000 CT-dependent host proteins found in diarrheal fluid are released from goblet cell granules along with the mucins that constitute the intestinal mucus layer. Our data also suggest that CT not only stimulates secretion or release of host proteins into the lumen, but that this potent toxin also impedes release of host factors. Several proteins belonging to protease inhibitor families were less abundant in both WT infection and after CT administration compared to V*. cholerae* Δ*ctx* infection (Fig S1). Thus, CT may increase the abundance of intestinal proteases, modifying proteolytic outcomes and thus the proteomic composition of choleric diarrhea.

The consequences of CT modulation of host factors implicated in innate defense, particularly on *V. cholerae*’s growth and survival in the intestinal niche, require further study. SP-D, one of the CT-dependent host factors we identified, was found to be a novel intestinal mucosal defense factor. This C-type lectin binds to L-glycero-D-mannoheptose (Hep), a constituent of the partially conserved lipopolysaccharide (LPS) inner core of many Gram-negative bacteria, including *V. cholerae* (43). SP-D was associated with the *V. cholerae* cell surface during infection and led to *V. cholerae* aggregation *in vitro*. Comparisons of *V. cholerae* growth in *sftpd*^+/+^ and *sftpd*^−/−^ infant mice, showed that SP-D protects against *V. cholerae* colonization, providing a new role for this lectin that has been linked to pulmonary defense against fungal, viral and bacterial pathogens (42,44,68,69). Strikingly, protection afforded by SP-D against *V. cholerae* colonization appeared to be restricted to the proximal portion of the small intestine. We previously observed a similar localized phenotype for *V. cholerae* colonization in infant mice lacking D-amino acid oxidase (DAO) (70), and propose that these observations reveal regional specificity to small intestinal mucosal defense factors. Although it seems paradoxical that *V. cholerae* would stimulate release of a factor such as SP-D that inhibits its own colonization, this may instead reflect the massive net gain that CT provides *V. cholerae* with respect to transmission. Thus, even if CT induces SP-D as part of a host defense program, *V. cholerae* still benefits from the toxin’s presence. It remains to be seen whether other enteric pathogens, including those that rely on secreted toxins for pathogenesis, also induce SP-D release, and whether this release is beneficial or antagonistic to the pathogen.

By honing our proteomic approach with SPLASH/M, we revealed the *in vivo V. cholerae* surface proteome as well as the complement of host proteins bound to the pathogen’s surface. Among the most abundant bacterial surface proteins during infection were TcpA and a methyl-accepting chemotaxis (VCA0176), two *V. cholerae* proteins that are known to be immunogenic (71), suggesting SPLASH/M-defined bacterial proteins represent antigenic targets that may be of therapeutic use.

SPLASH/M also enabled the unbiased identification of a class of host proteins we termed <u>h</u>ost-derived <u>b</u>acterial <u>b</u>inding <u>p</u>roteins (HBBPs) that were more abundant on the *V. cholerae* surface than in diarrheal fluid. This included Intelectin, which is known *V. cholerae-* targeting HBBPs implicated in microbial recognition that were identified with different methods (32,72). Only ~25% of the identified HBBPs have previously been linked to host defense/ inflammation. Most of the other HBBPs were classified as enzymes or linked to metabolism, raising the possibility that these factors might impact the *in vivo* physiology of *V. cholerae* and other microbes through binding; alternatively, the apparent binding of some of these factors to the *V*. *cholerae* surface could be fortuitous.

Three HBBP chosen for additional analysis, AnxA1, LPO and ZAG, bound to *V. cholerae* cells grown in laboratory media without additional host factors present (Fig. 4), providing evidence that they can directly bind the pathogen. Although only ZAG and LPO bound to specific and distinct microbial glycans, since AnxA1 belongs to the annexin superfamily of calcium-dependent phospholipid-binding-proteins, it may instead bind a non-carbohydrate ligand on the bacterial cell surface, such as phosphatidylserine (PS), a lipid constituent of the bacterial membrane (54). We found that 1-10% of fecal microbiota were bound by AnxA1, ZAG and LPO, a similar range as reported for IgA and SP-D (39,63). Each of these 3 proteins bound to distinct microbial taxa, raising the possibilities that these proteins and other HBBPs might play a role in general host microbial surveillance and modify the composition and/or function of the intestinal microbiome. These SPLASH/M-identified interactions may not necessarily be antagonistic, and could have important and far-reaching consequences on host physiology. For example, mice deficient in SP-D have distinct gut microbiota and immune profiles (39).

While our use of the infant rabbit model of cholera facilitated the development of SPLASH/M, this approach should be applicable to additional pathogenic and non-pathogenic microbes alike. Methods to isolate particular microorganisms, such as FACS, will facilitate SPLASH/M-based definitions of the *in vivo* proteomic landscapes of microbe-host interfaces. Furthermore, variants of this approach should be applicable to reveal this interface for microbes that grow intracellularly (6–9). Additional efforts to reveal the full complement of HBBPs during infection with different pathogens in different tissues as well as their functions will reveal valuable new insights into microbial and host biology. Moreover, since gut symbionts are intrinsically linked to host physiology and health, investigation of HBBP coating of symbionts in different contexts, such as obesity, will offer new perspectives in pathophysiology. Ultimately, defining the proteomic composition of the microbe-host interface will deepen our understanding of interkingdom interactions that underlie homeostasis and disease, and offer new factors to target for therapeutic applications.

## Acknowledgments

We thank members of the Waldor research group for helpful discussions. Alyson R. Warr for expert help with bioinformatics. Sharon Prentice for technical assistance and the Bettencourt-Schueller foundation for support. Glycomic experiments were done with the participation of the Protein-Glycan Interaction Resource of the CFG, and the National Center for Functional Glycomics, supporting grant P41 GM103694 and R24 GM137763. We thank the ThermoFisher Center for Multiplexed proteomics of Harvard Medical School for isobaric tandem mass tag proteomic. Work in M.K.W laboratory is supported by HHMI and NIH grant R01 AI-042347. T.Z. was supported by a Sarah Elizabeth O’Brien Trust Postdoctoral Fellowship. A.Z. was supported by an EMBO long-term fellowship (ALTF 1514-2016) and by a HHMI Fellowship of the Life Sciences Research Foundation.

## Authors contribution

A.Z. and M.K.W. conceived and designed the study. A.Z., H.Z., B.F., and C.J.K, performed all experiments; A.Z. and R.T.G., analyzed data. A.Z. and M.K.W. wrote the manuscript and all authors edited the paper.

**Figure S1:**
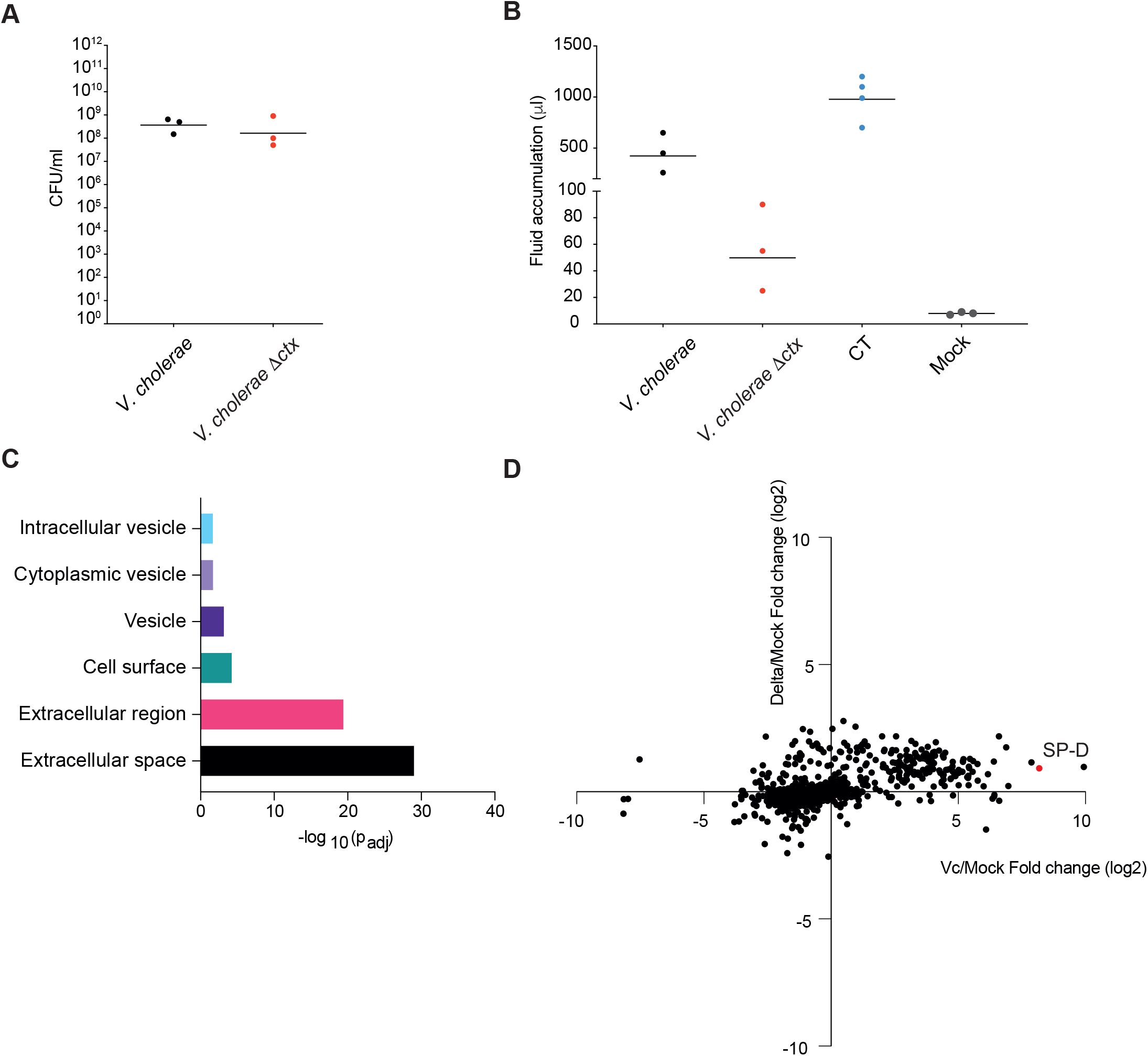
Diarrheal fluid proteomic response to V. cholerae is largely driven by CT. (A) Bacterial burdens recovered from diarrheal fluid harvested from *V. cholerae* and *V. cholerae* Δ*ctx* infected rabbits. (B) Diarrheal fluid volumes collected from rabbits infected with *V. cholerae*, *V. cholerae* Δ*ctx*, purified cholera toxin (CT) and buffer (Mock). (C) Predicted localization of rabbit proteins identified in diarrheal fluid. Bioinformatic analysis was performed using the G:Profiler (http://biit.cs.ut.ee/gprofiler/) webtool for finding enriched GOcatgories. (D) Scatterplot of relative fold changes in protein abundances isolated from rabbit infected with *V. cholerae* Δ*ctx* (Delta) compared to wild-type *V. cholerae* (Vc), each relative to the proteomes of mock infected animals. The red dot indicates SP-D.

**Figure S2:**
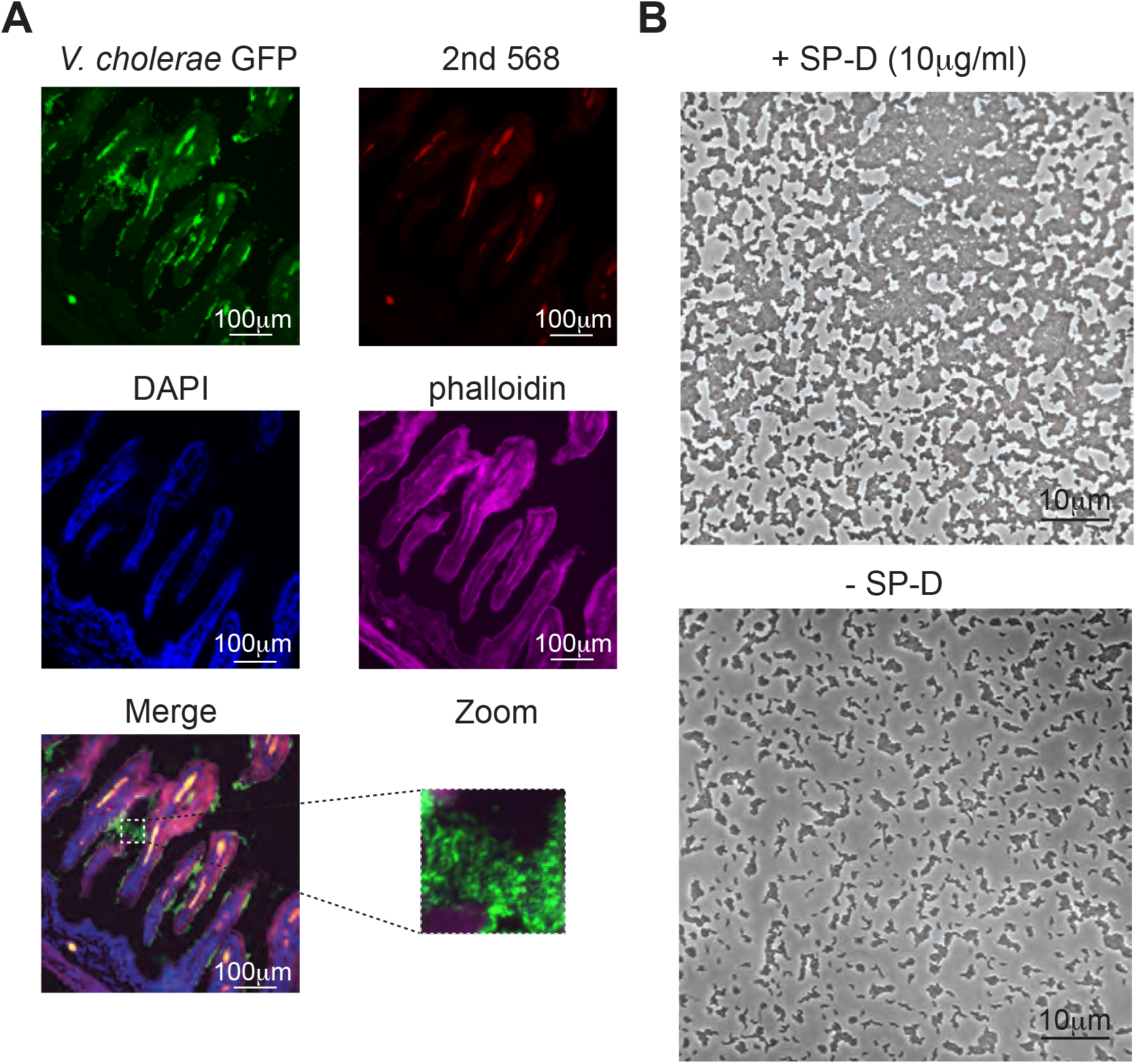
SP-D is an intestinal mucosal defense factor. (A) Immunofluorescence micrographs of rabbit small intestines inoculated with GFP-expressing *V. cholerae*. Bacterial cells were detected by GFP fluorescence. Phalloidin (for actin labeling) is stained with an antibody coupled to Alexa fluor 647 and DAPI (for DNA labeling) is shown is blue. Only anti-goat antibody coupled to Alexa fluor 468 was used to assess unspecific staining of the second antibody. Scale bar is 100 μm. (B) Wide field of micrographs shown in Fig. 2C. Scale bar is 10 μm.

**Figure S3:**
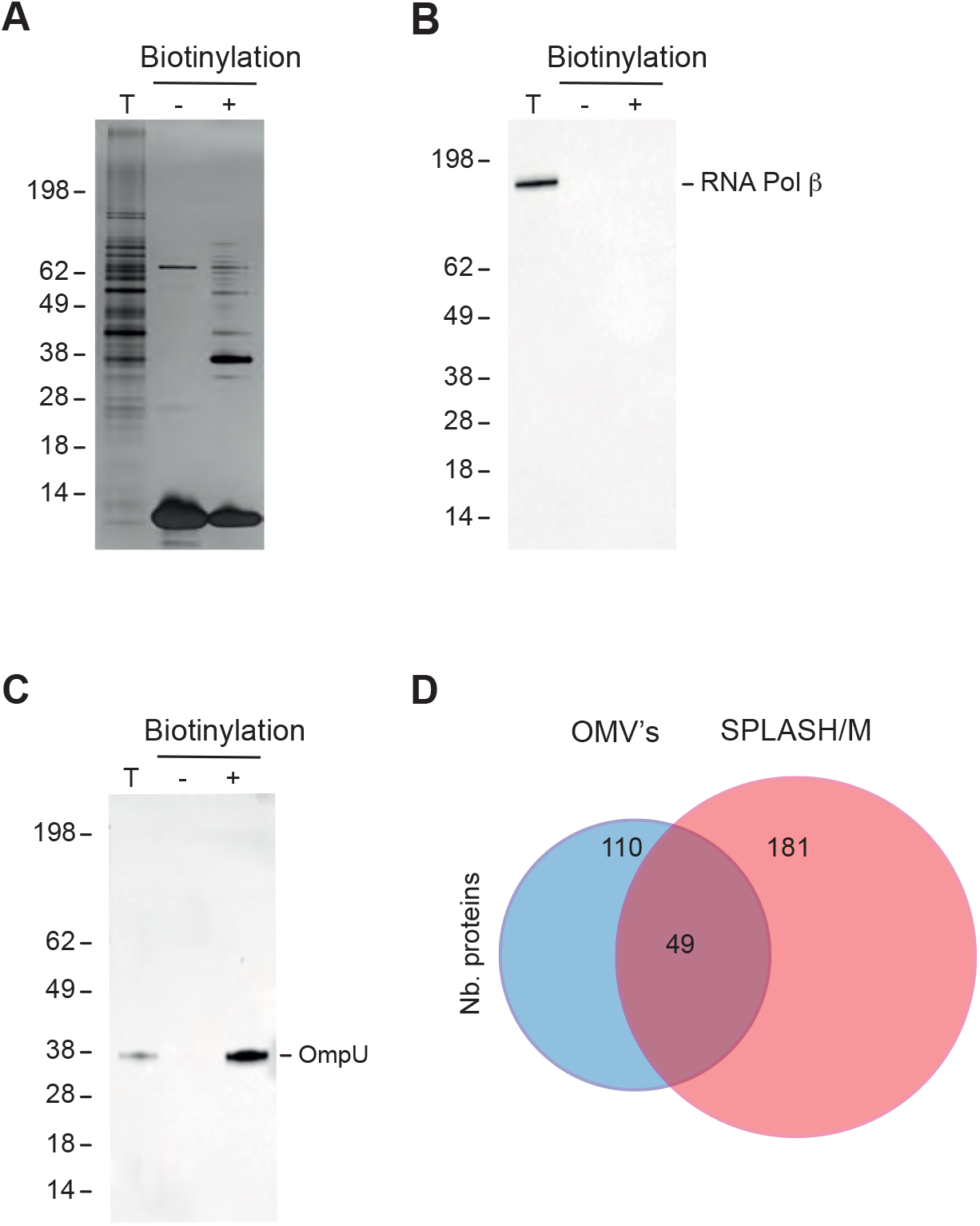
Identification of surface-exposed *V. cholerae* proteins and *V. cholerae*-bound host-derived proteins. Controls validating surface biotinylation for SPLASH/M (A-C). Proteins isolated following SLASH/M protocol with (+) or without (-) the biotinylation step were separated by 10% acrylamide SDS-PAGE and silver-stained (A). Presence of cytoplasmic RNA polymerase β (B) and outer-membrane OmpU (C) were assessed by immunostaining with anti-RNApol and anti-OmpU antibodies, respectively. T: total *V. cholerae* lysate. (D) Venn diagram showing the comparison of *V. cholerae* proteins identified with SLASH/M and *V. cholerae* outer membrane vesicles (OMV’s) proteomes; 181 and 110 are the total number of proteins from each group (31).

**Figure S4:**
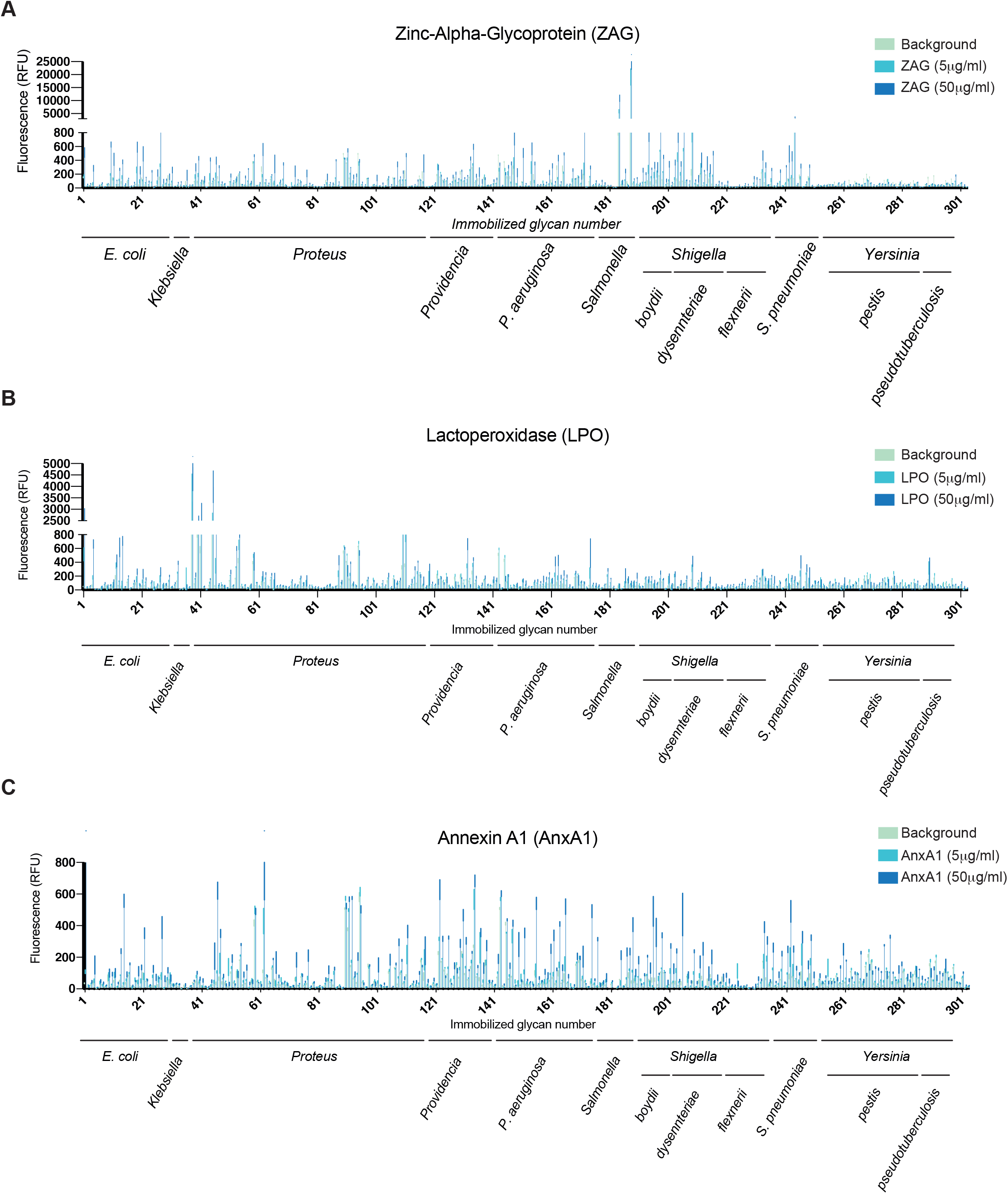
LPO, AnxA1 and ZAG binding to microbial glycans. (A-C) Results of ZAG (A), LPO (B) and AnxA1 (C) binding to Microbial Glycan Microarray organized by genus and species. Data are presented as the mean ± s.d. (n=4 of a technical replicate for each immobilized glycan). Note: scales on Y axes are different. The complete datasets are available in Supplementary Table S4.

**Figure S5:**
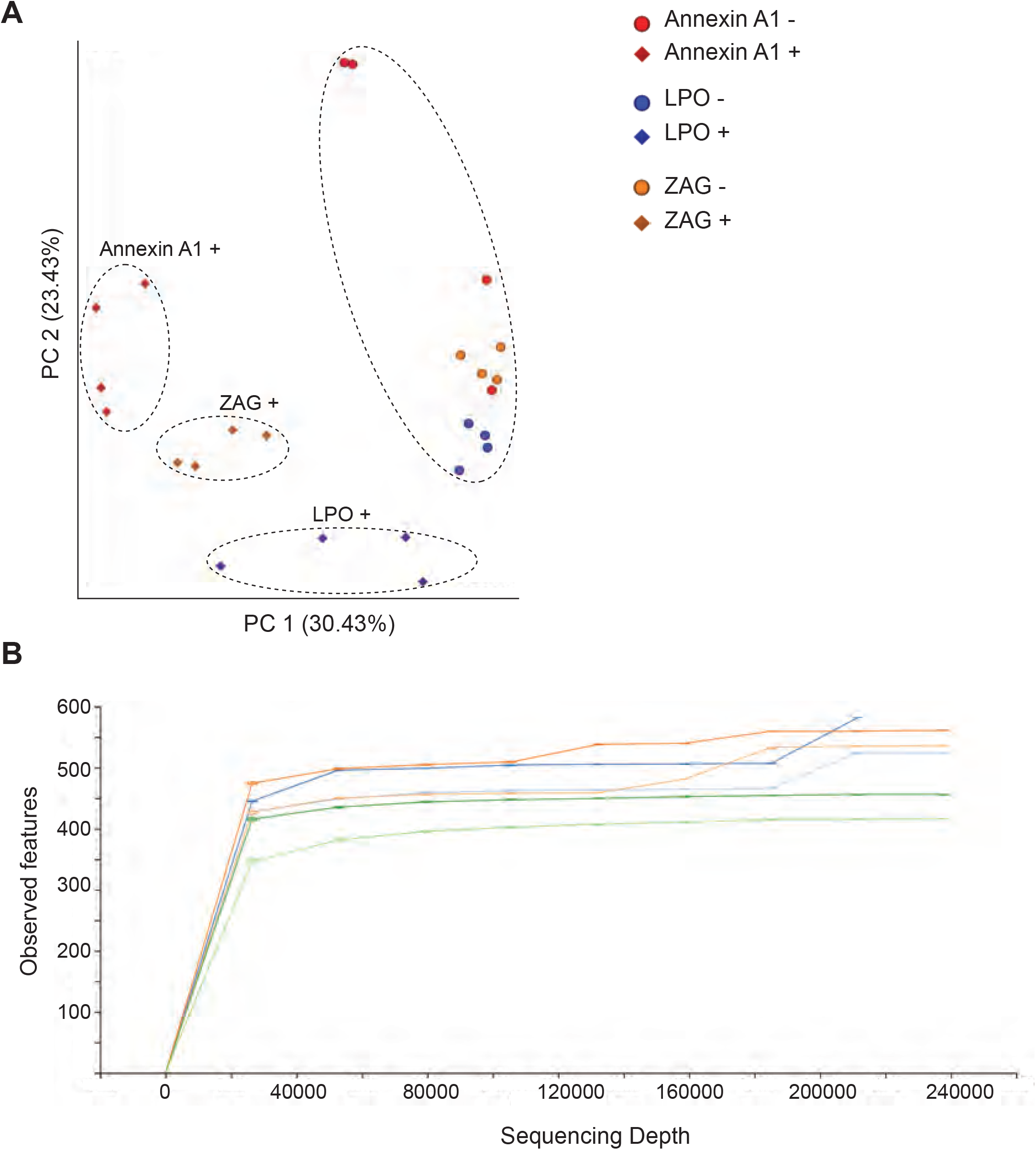
HBBPs interact with gut commensal bacteria. (A) Principle coordinate analyzes based on the Bray Curtis β-diversity metric showing that samples for each AnxA1, LPO or ZAG positive population cluster together while all the HBBP negative populations cluster together. (B) Alpha rarefaction plot. Shown are the number of different observed features as a function of the number of sequences analyzed and generated with QIIME2.

## Materials and Methods

### Ethics Statement

Animal experiments were conducted according to protocols approved by the Brigham and Women’s Hospital Committee on Animals (Institutional Animal Care and Use Committee protocol number 2016N000334 and Animal Welfare Assurance of Compliance number A4752-01) and in accordance with recommendations in the National Institute of Health’s Guide for the Care and Use of Laboratory Animals and the Animal Welfare Act of the United States Department of Agriculture.

### Bacterial strains, and growth condition

*V. cholerae* strain H1, a clinical isolate from 2010 and its Δ*ctx* derivative (34,65) were cultured in Luria-Bertani (LB) medium or on LB agar plates at 37°C unless otherwise stated, supplemented with streptomycin at a concentration of 200 μg/ml. *V. cholerae* cells carrying the pUA-GFP plasmid, which contains a GFP gene under strong constitutive promoter) (73) was used for immunostaining of infected infant rabbit small intestine and cultured overnight at 30°C in LB supplemented with streptomycin (200 μg/ml) and kanamycin (50 μg/ml).

### Infant rabbit infection studies

For inocula preparation, overnight bacterial cultures were diluted 1:100 in 50 mL LB and cultured with aeration at 37°C until OD600 0.5-0.9. ~2×10^10^ CFU were pelleted by centrifugation at 5000 g for 5 min, the supernatant was removed, and cell pellets were re-suspended in 10 mL of 2.5% sodium bicarbonate solution (2.5g in 100 mL water; pH 9.0) to a final cell density of ~2×10^9^ CFU/ml. Serial dilutions of the inoculum were plated to enumerate the inoculum dose. Infant rabbit infections were performed as previously described (24,28). Briefly, two-day old litters of mixed gender New Zealand White rabbit were co-housed with a lactating dam (Charles River) for the duration of the experiment. Each infant rabbit was orogastrically inoculated with 500 μl of the inoculum, using a size 4 French catheter. Following inoculation, the infant rabbits were monitored at least 2x/day for signs of illness and euthanized ~16-18 hours post infection. For purified cholera toxin (CT) experiments, 50 μg CT (Sigma, C8052) was used per rabbit (500 μl of a 100 μg/ml solution in sodium bicarbonate). Animals infected with CT were euthanized 3-6 h post-inoculation.

### Mice colonization assay

C57BL/6 Sftpd^−^/^−^ mice were purchased from Jackson laboratory and were bred at the Harvard Institutes of Medicine animal facility. Littermates that were the offspring of heterozygous Sftpd^+^/^−^ breeders were used in this study. Infant mice were genotyped post-mortem at the end of the colonization assay. Intestinal colonization in infant mice was conducted as described (74). Briefly, bacterial cells were grown overnight at 30°C and then diluted 1:1000 in LB. Infant mice were orogastrically inoculated with 50 μl (~ 10^5^ cfu) and then sacrificed after ~ 18 hours. Small intestines were equally divided into proximal and distal segments. Dilutions of small intestines homogenates were plated on LB agar plates supplemented with 200 μg/ml streptomycin to enumerate CFU. Statistical significance was determined using a Mann-Whitney U t test. Infant mice were genotyped post-mortem at the end of the colonization assay using tail chips and PCR according to Jackson laboratory protocol using primers 24516 (TGT TGA TGC ATG TTA TGT GAT GA), 24517 (CCT AGG GAA GGC TAG GGA GT) and oIMR2088 (AGA CTG CCT TGG GAA AAG CG).

### Immunofluorescence microscopy

Immunofluorescence images were analyzed from 6 rabbits infected with *V. cholerae*-GFP; 2 or 3 sections of the small intestine per rabbit were examined. Briefly, tissue samples used for immunofluorescence were fixed in 4% PFA for 2 hours, and subsequently stored in 30% sucrose prior to embedding in a 1:2.5 mixture of OCT (Tissue-Tek) and stored at –80°C, as previously described (75). Frozen sections were then cut at a thickness of 10-15 μm using a cryotome (catalog no. CM1860UV; Leica). Sections were first blocked with 5% bovine serum albumin (BSA) in PBS for 1 h and then stained overnight at 4°C with a primary anti-SP-D antibody (1:500, R&D Systems, AF1920), diluted in PBS with 0.5% BSA and 0.5% Triton X-100, anti-GFP labeled with Alexa 488 (1/1,000, SAB4600051). After washing 3x with 1× PBS containing 0.5% Triton X-100, sections were incubated with Alexa Fluor 647 phalloidin (1/1000; Invitrogen) and anti-Goat Alexa Fluor 568 (1/1000, ThermoFisher, A-11055) for 1◻h at room temperature, washed, and stained for 5◻min with 4′,6-diamidino-2-phenylindole (DAPI) at 2◻μg/ml for 10◻min, and covered with ProLong Diamond mounting medium. Following staining, slides were imaged using a Nikon Ti Eclipse equipped with a metal-oxide-semiconductor (sCMOS) camera (Andor Zyla) for wide-field microscopy.

### Preparation of diarrheal fluid for MS analysis and immunoblotting

Diarrheal fluids were filtered through sterile polyester membranes with a pore size of 0.22 μm before precipitation with trichloroacetic acid (TCA) 15%, 45 min on ice. Precipitated proteins were wash once in acetone and resuspended in 1X blue loading buffer (NEB, B7703S).

For immunoblotting, bacterial pellets or precipitated proteins were resuspend in blue loading buffer (NEB, B7703S), boiled at 95°C for 10 min and loaded on 10% gels (Bio-Rad) for electrophoresis. Proteins were transferred from the gel to nitrocellulose membranes and immunoblotted. Antibodies for western blot assays were used at the following concentrations: anti-SP-D (1:2,000, R&D Systems, AF1920), anti-LPO (1:2000, LSBio, LSl1JC25068), anti-AnxA1 (1:500, ThermoFisher, 71-3400), anti-ZAG (1:2000, ThermoFisher, H00000563-B01P), anti-RNA Polymerase (1:2000, Biolegend, 663903) and anti-OmpU (1:500, homemade, gift from the Mekalanos lab). The membranes were developed with SuperSignal West Femto maximum-sensitivity substrate (ThermoFisher) and visualized with a ChemiDoc Scientific imaging system (BioRad).

### Peptide Labeling with Tandem Mass Tags and Mass Spectrometry

Samples were submitted in 1X blue loading buffer (NEB, B7703S) to the Thermo Fisher Center for Multiplexed Proteomics at Harvard Medical School (Boston, MA, USA) for Isobaric Tandem Mass Tag (TMT)-based quantitative proteomics. Briefly, after adjusting proteins to equal concentrations, 40 μl of each sample was loaded on 10% Bis/Tris gels and run at 120V for 10 min in MES buffer. Gel bands were cut out, destained, reduced and alkylated. In-gel Trypsin digests were performed overnight and peptides were extracted and labeled with TMT10 reagents. Labeling reactions were combined, cleaned, and dried down. Peptides were resuspended in 5% Acetonitrile, 5% formic acid and 1/3 of the sample was shot on an Orbitrap Fusion Mass spectrometer. Peptides were detected (MS1) and quantified (MS3) in the Orbitrap Fusion Mass spectrometer. Peptides were sequenced (MS2) in the ion trap. MS2 spectra were searched using the SEQUEST algorithm against a Uniprot composite database derived from the combined *V. cholerae* and *Oryctolagus cuniculus* (rabbit) proteomes containing its reversed complement and known contaminants. Peptide spectral matches were filtered to a 1% false discovery rate (FDR) using the target-decoy strategy combined with linear discriminant analysis. Proteins were quantified only from peptides with a summed signal/noise (SN) threshold of >=200 and MS2 isolation specificity of 0.5.

### Gene set enrichment analysis

The G:Profiler (http://biit.cs.ut.ee/gprofiler/) webtool was used for finding enriched GO cellular component terms in the rabbit intestinal proteome. A score above 1.8 for negative log of adjusted p-values was considered significant. Gene set enrichment was performed as previously described (76) using fast GSEA (fGSEA) in R (version 1.8.0) (77) with modifications. Only genes with annotation were considered. The normalized mean proportion for each protein was divided by the value of that protein in the uninfected data set and Log2 transformed to create a fold change. These Log2 fold change values were use as the “rank” for fGSEA.

### Hierarchical clustering

The SN of each protein was first normalized by calculating the proportion of the total signal represented by that protein in a given sample. Clustering was then performed on these values in R using heatmap.2 with the default Pearson correlation method.

### *In vitro* protein-V. cholerae binding assay

Binding assays were carried out as previously described (32). Briefly, bacteria were grown to O.D ~ 0.4 in LB and then centrifuged (5,000 *g*, 5min at RT). Bacterial pellets were washed twice in 25 ml HEPES-buffered saline (140 mM NaCl, 1.5 mM Na_2_HPO_4_, 50 mM HEPES, pH 7.5) supplemented with 5 mM CaCl_2_. Bacterial cells were then incubated with 0.25 μg of purified human SP-D (R&D Systems, 1920-SP-050), ZAG (R&D Systems, 4764-ZA-050), AnxA1 (R&D Systems, 3770-AN-050) or LPO (MyBiosource, MBS954610) for 30 min at RT and washed twice with an equal volume of buffer. Bacterial pellets were then resuspended in 1X blue loading buffer (NEB, B7703S). Unbound input and the two washes were treated with 4X blue loading buffer (NEB, B7703S) and incubated at 95°C for 10 min prior to SDS-PAGE and western blot analysis. All binding experiments were repeated at least 3x with consistent results.

### Bacterial aggregation assay

*V. cholerae* cells were grown to O.D ~ 0.4 in LB, centrifuged (5,000 *g*, 5 min at RT) and then resuspended and washed in phosphate-buffered saline (PBS) supplemented with 5 mM CaCl_2_. Bacterial suspensions were incubated with human SP-D at a concentration of 10mg/ml (R&D Systems, 1920-SP-050) for 1h at RT without agitation and observed by light microscopy (Nikon Ti Eclipse equipped with a metal-oxide-semiconductor (sCMOS) camera (Andor Zyla)). Figures were made using Fiji software (version 2.1.0/1.53c).

### Surface Protein LAbelingS Host/Microbe (SPLASH/M)

Diarrheal fluid was harvested 16-18 hr after inoculation of infant rabbits. The fluid was then filtered through a 5 μM filter to remove particulate matter and eukaryotic cells. Bacteria were isolated from diarrheal fluid by centrifugation (5000 × *g*, 5 min at RT). Bacterial pellets were washed twice and resuspended in phosphate-buffered saline (PBS) supplemented with 1 mM CaCl_2_, 0.5 mM MgCl_2_ and 1.5 mM D-biotin at RT. Cell surface biotinylation were performed as described (ref) with modifications. Sulfo-NHS-LC-biotin (ThermoFisher, 21335) was added to a final concentration of 200 μM for 20 min at RT. The reaction was stopped by addition of 2 volumes of buffer (80mM Tris pH7, 100mM NaCl, 30mM KCl, 1mM CaCl_2_ and 0.5mM MgCl_2_). After washing the bacterial cells 3x with the same buffer they were resuspended in 50 mM Tris pH 7, 50 mM NaCl, 10 mM MgCl_2_, DNAse (0.1mg/ml), lysozyme (0.1mg/ml) and Complete protease inhibitor mixture (Roche). Cells were broken using an Emulsiflex-C3 (Avestin) and the crude membrane fraction was isolated by ultracentrifugation at 45,000 g for 45 min. Membrane-containing fractions were washed twice in 50 mM Tris pH 7, 150 mM KCl, 10 mM EDTA and Complete protease inhibitor mixture (Roche). Membranes were then solubilized overnight at 4°C in presence of 0.5% n-Dodecyl β-D-maltoside (DDM) (Sigma, D4641). Lysates were used for co-immunoprecipitation using Dynabeads M-280 Streptavidin (ThermoFisher, 11205D) overnight at 4°C. Magnetic beads were washed thrice with 1 ml of Tris pH 7, 100 mM NaCl and 0.2% Tween 20 and resuspended in 50 μL of 1X blue loading buffer (NEB, B7703S) and heated for 10 min at 96°C.

### HHBP binding to glycan arrays

Human ZAG (R&D Systems, 4764-ZA-050), human LPO (MyBiosource, MBS954610) and human AnxA1 (R&D Systems, 3770-AN-050) were provided to the Protein-Glycan Interaction Resource at the National Center for Functional Glycomics (Beth Israel Deaconess Hospital, Boston) for hybridization to the Microbial Glycan Microarray (MGM). The MGM array were prepared as previously described (62). The printed array includes polysaccharides derived from 313 different bacteria printed at 500 μg/ml, in replicates of 6. To interrogate the MGM, ZAG, LPO and AnxA1 were diluted to 5 μg/ml and 50 μg/ml in binding buffer (20 mM Tris-HCl, pH 7.4, 150 mM NaCl, 2 mM CaCl_2_, 2 mM magnesium chloride (MgCl_2_), 1% BSA and 0.05% Tween 20) and applied directly to the array surface for 1 h. After incubation, the array was washed by soaking with binding buffer four times. ZAG was detected with Anti-ZAG (2 μg/ml, AZGP1 antibody, H00000563-B01P MaxPab) and anti-mouse IgG-Alexa-488 (5 μg/ml) diluted in binding buffer, applied directly to the array surface and allowed to incubate for 1 h. Similarly, LPO was detected with Anti-LPO (2 μg/ml, Thermofisher PA5-18917) and anti-goat IgG-Alexa-488 (5 μg/ml) diluted in binding buffer, and then applied directly to the array surface for 1 h. Similarly, AnxA1 was detected with Anti-AnxA1 (2 μg/ml, Sigma, AMAB90558) and anti-mouse IgG-Alexa-488 (5 μg/ml). The arrays were washed in binding buffer (four times), binding buffer without BSA and Tween 20 (four times) followed by deionized water (four times) and scanned. The high and low fluorescence values from the six replicates were eliminated and the remaining four values were averaged. Data were plotted with Excel (Microsoft) as average relative fluorescence units (RFU) versus print identification number. The top 5 HBBP-glycans interactions for ZAG and LPO showed in Fig. 4GH were defined as 3-fold over background and exhibiting a dose-responsive binding which was not the case for AnxA1.

### Analysis of HBBPs binding microbiota

Fecal pellets from specific pathogen free (SPF) C57BL/6 mice were collected and directly resuspended in phosphate-buffered saline (PBS) (100 mg of feces in 100 μl) supplemented with 1% BSA and 1 mM CaCl_2_, and filtered with a 40 μm cell strainer to remove particulate matter.

Bacterial suspensions were centrifuged (5,000 g, 5min) and washed twice in the same buffer. 20 μl of bacterial suspension was incubated with 2 μg of biotinylated anti-LPO (LSBio, LS-C684314), anti-ZAG (R&D Systems, BAF4764) or anti-AnxA1 (LSBio, LS-C317217) for 30 min on ice. After washing 3x and resuspension in PBS, supplemented with 1% BSA and 1 mM CaCl_2_, bacteria were incubated with 1 μg of Streptavidin-PE-Cy7 (ThermoFisher, SA1012) for 15 min on ice. After washing, bacterial genomic DNA was stained with 1/10 000 dilution of SybrGreen followed by two washes. Bacterial suspensions were then analyzed by flow cytometry (Sony, SH800) and HBBP-positive or HBBP-negative population were sorted.

### 16s rRNA gene sequencing and analysis

PureLink Microbiome DNA Purification Kit (ThermoFisher, A29790) was used according to the manufacture protocol to extract the DNA from the sorted-microbiota. 16s rRNA amplification were done as previously described (70). Briefly, the V1–V2 region of 16S ribosomal RNA was PCR amplified (12.5 ng purified DNA per reaction; Phusion polymerase, New England Biolab) for 25 cycles (95°C for 30s, 50°C for 30s and 72°C for 30s) (primer pair: 27Fmod/338R (Ref)). PCR products were then purified (MinElute, QIAGEN) and resuspended in 25μl of 10mM Tris-HCl pH8.5. The V1–V2 PCR products were indexed with the Nextera XT Index kit (Illumina) by PCR (2.5μl PCR product; Nextera XT Index primers; Phusion polymerase) for eight cycles (95°C for 30s, 55°C for 30s, and 72°C for 30s). The 16S rRNA amplicons with indices were purified (MinElute, QIAGEN), resuspended in 25μl of 10mM Tris-HCl pH8.5, quantified with a Qubit 2.0 Fluorometer (Life Technologies), pooled at a concentration of 4nM, denatured, diluted to a final concentration of 4pM and sequenced using the MiSeq Reagent Kit v3 (600-cycle, paired-end, Illumina) on a MiSeq sequencer (Illumina). Sequencing reads were demultiplexed using MiSeq Reporter v2.0 and further processed using QIIME2 (Ref). Briefly, paired-end reads (FASTQ files) were merged with FastqJoin and quality filtered with a Q-score cutoff of 20. Merged sequencing reads were denoised using using DADA2 (Ref). Taxonomic classification was generated using a pre-trained naive Bayes classifier on the basis of the bacterial 16S rRNA Greengenes reference database and QIIME2 (https://qiime2.org).

## CONTACT FOR REAGENT AND RESOURCE SHARING

Further information and requests for resources and reagents should be directed to and will be fulfilled by the Lead Contact, Matthew K Waldor

## Supplementary items

Table S1: Diarrheal fluid proteomes

Table S2: Pathway enrichment

Table S3: SPLASHM proteome

Table S4: Glycomic data

